# Diffusion MRI of white matter microstructure development in childhood and adolescence: Methods, challenges and progress

**DOI:** 10.1101/153924

**Authors:** Christian K. Tamnes, David R. Roalf, Anne-Lise Goddings, Catherine Lebel

## Abstract

Diffusion magnetic resonance imaging (dMRI) continues to grow in popularity as a useful neuroimaging method to study brain development, and longitudinal studies that track the same individuals over time are emerging. Over the last decade, seminal work using dMRI has provided new insights into the development of brain white matter (WM) microstructure, connections and networks throughout childhood and adolescence. This review provides an introduction to dMRI, both diffusion tensor imaging (DTI) and other dMRI models, as well as common acquisition and analysis approaches. We highlight the difficulties associated with ascribing these imaging measurements and their changes over time to specific underlying cellular and molecular events. We also discuss selected methodological challenges that are of particular relevance for studies of development, including critical choices related to image acquisition, image analysis, quality control assessment, and the within-subject and longitudinal reliability of dMRI measurements. Next, we review the exciting progress in the characterization and understanding of brain development that has resulted from dMRI studies in childhood and adolescence, including brief overviews and discussions of studies focusing on sex and individual differences. Finally, we outline future directions that will be beneficial to the field.

## 1. Introduction

Longitudinal magnetic resonance imaging (MRI) studies provide evidence for substantial developmental macrostructural changes in the brain continuing throughout childhood and adolescence. These changes are tissue specific, and involve decreases in gray matter (GM) volume and increases in white matter (WM) volume (Brain Development Cooperative Group, 2012; Giedd et al., 2015; Lebel & Beaulieu, 2011; Mills et al., 2016). The biophysical mechanisms of WM changes in this period of life are multiple, but include further axon myelination via thickening of the myelin sheets and axonal growth from increasing caliber (Paus, 2010). Diffusion MRI (dMRI) is a method that allows for further understanding of the living human brain and how it develops, especially for WM anatomy, as it yields quantitative parameters related to tissue microstructure (Tournier et al., 2011). Hence, dMRI is a promising technique that may bring neuroimaging studies one step closer to the underlying cellular and molecular processes.

In the current review, we will describe and discuss dMRI methods available to study the developing human brain and review the recent progress made using these increasingly popular methods. First, we will provide an introduction to dMRI, including overviews of different acquisition and analysis approaches. Throughout this section we will discuss selected methodological challenges, focusing on issues of particular importance for developmental studies. Next, we will review dMRI studies of WM microstructure development in childhood and adolescence, emphasizing longitudinal studies where available. Finally, we will outline selected future directions that we believe will be beneficial to the field.

## 2. Diffusion MRI: acquisition and analysis

### 2.1. Modelling diffusion

dMRI exploits a phenomenon that is a nuisance in many MRI sequences. Specifically, the random movement of water in the brain reduces the measurable MRI signal. While this phenomenon reduces the signal for other modalities (e.g., T1, T2), dMRI uses it to measure diffusivity and obtain information about the underlying tissue microstructure. Using diffusion sensitizing gradients and non-diffusion weighted references image(s), one can obtain estimates of the apparent diffusion coefficient (ADC) in one or more directions. Barriers such as cell membranes and myelin prevent diffusion from occurring freely in the human brain, and diffusion coefficients vary at different locations and orientations. This is particularly true of WM, which is highly ordered and has higher water diffusivity parallel to rather than perpendicular to nerve fibers (Chenevert et al., 1990; Doran et al., 1990; Douek et al., 1991). Given the complex organization of the brain, ADC must be measured in multiple directions to fully characterize *in vivo* diffusion.

Typical quantification of dMRI is achieved in a tensor model. The tensor is a mathematical construct that can be used to characterize diffusion in all directions, and this can be calculated by collecting data using at least 6 non-collinear diffusion gradient directions; this is called diffusion tensor imaging (DTI, see (Mori & Tournier, 2013) for an introduction). The tensor is diagonalized to calculate three eigenvector/eigenvalue pairs which represent the direction and magnitude of diffusivity along the three orthogonal axes (*v*_1_, λ_1;_ *v*_2_, λ_2_; *v*_3_, λ_3_). The vectors represent the primary, secondary and tertiary diffusion directions, respectively, with the primary diffusion direction generally assumed to correspond to the primary fiber direction within a voxel. The eigenvalues can be used to quantify diffusion properties in several ways. Most commonly, fractional anisotropy (FA) is used as a measure of the directionality of diffusion. FA is essentially the variance of the eigenvalues (between 0-1), with 1 being highly directional diffusion and 0 being completely isotropic diffusion. Mean diffusivity (MD) is the mean of the 3 eigenvalues and represents the average magnitude of diffusion; axial diffusivity (AD; also called parallel or longitudinal) is diffusivity along the longest axis of the ellipsoid (λ_1_); radial diffusivity (RD; also called perpendicular or transverse) is the average of the diffusivity values along the two minor axes of the ellipsoid (λ_2_, λ_3_). These parameters vary across individuals, and regionally within the brain. They are also dependent on the hardware, software, acquisition parameters, processing, and analysis methods used, which is important to keep in mind during any comparisons across studies.

Diffusivity is influenced by many aspects of brain micro- and macro-structure including myelin content, cell density, axon diameter, axon coherence, membrane permeability, and water content (Beaulieu, 2002). During healthy brain development, changes to diffusion parameters are generally attributed to normal maturation processes such as myelination and increased axonal size and packing. AD and RD provide slightly higher specificity compared to FA and MD: animal models have shown that RD is altered by demyelination and remyelination while AD remains unchanged by myelin changes, but is altered by axonal injury (Song et al., 2003; Song et al., 2002; Song et al., 2005). Axonal injury is not expected to occur during normal brain development, but AD changes have been linked to straightening of axons, which does occur during development (Takahashi et al., 2000). While diffusion parameters are sensitive to these changes, they are not specific to them, and one must remember, as Jones et al., (2013) state in a recent commentary, diffusion imaging only measures one thing – “the dephasing of spins of protons in the presence of a spatially-varying magnetic field” (p.239). While dephasing of spins provides information about the microstructural organization of the brain, it is incomplete information and mapping the outcome of complex diffusion analyses onto specific microstructural traits (e.g., axonal diameter, myelination) is challenging, if not impossible.

DTI is by far the most common method used to date to characterize WM changes in neurodevelopment, but it has numerous limitations and interpretation of its parameters is not necessarily straightforward (Jones & Cercignani, 2010; Tournier et al., 2011). Only one fiber direction per voxel can be modelled in DTI, yet as many as 60-90% of WM voxels in the brain contain multiple fiber populations (Jeurissen et al., 2013). High angular resolution diffusion imaging (HARDI) is an advanced dMRI acquisition technique where many more diffusion encoding gradient directions are measured compared to DTI. This enables more advanced modelling of the diffusion signal within each voxel, for example using diffusion spectrum imaging (DSI) (Wedeen et al., 2005) or q-ball imaging (Tuch, 2004). However, HARDI methods have not been widely used to study neurodevelopment (yet), primarily due to two problems. First, they generally require long acquisition times that are not necessarily feasible in pediatric and adolescent populations. Second, it is difficult to identify an easily quantifiable parameter that can be compared across age to measure development. However, with hardware and software developments, acquisition times will become faster. Progress is also being made toward quantifiable and more interpretable measures and statistical methods that further probe WM microstructure (Raffelt et al., 2012; Raffelt et al., 2015). For instance, Raffelt et al. (2015) has coined the term fixel to refer to a specific population of fibers within a single voxel and proposed a method to perform whole-brain fixel-based analysis using probabilistic tractography, which shows promise for better handling crossing fibers and resolving tracts. Thus, in the coming years, these advanced methods may provide substantial insight into brain development patterns across childhood and adolescence.

Other recent developments in dMRI methods include diffusion kurtosis imaging (DKI) and neurite orientation dispersion and density imaging (NODDI). DKI is a method that accounts for the non-Gaussian signal decay that occurs due to restricted diffusion (Jensen & Helpern, 2010; Jensen et al., 2005), and produces measures of mean, axial, and radial kurtosis (analogous to mean, axial and radial diffusivity). NODDI (Tariq et al., 2016; Zhang et al., 2012) models tissue compartments for intracellular, extra-cellular, and cerebrospinal fluid (CSF) separately, and estimates both an orientation dispersion index (ODI) and neurite density index (NDI). Initial applications of these methods demonstrate utility in studying brain development (see section 3.1), and may become more common in future studies. However, as with HARDI methods, DKI and NODDI require a substantial increase in acquisition time compared to DTI: a limitation that is challenging in children.

### 2.2. Diffusion image acquisition

Data quality is one of the most important considerations when designing an acquisition protocol. Signal-to-noise ratio (SNR) is a measure of the average available signal compared to the typical background noise in an image. Sufficiently high SNR is necessary to allow robust measurement of diffusion parameters (FA, MD, etc.) and accurate assessment of images. SNR depends on both hardware (e.g., field strength) and acquisition protocol (e.g., echo time, number of averages, image resolution). SNR is tightly linked with scan time, and longer acquisitions permit higher SNR. Thus, all parameter choices for a given diffusion imaging acquisition sequence should be made while considering SNR to maximize image quality, while balancing time constraints to maximize participant compliance. However, recent advances in the use of multi-band or simultaneous multi-slice acquisitions (as e.g., used in the Human Connectome Project (HCP) Lifespan Pilot) may reduce scan time and allow for more complex multi-shell protocols, also in developmental samples. Furthermore, studies show that using sequences with phase encoding reversal (combining blip-up and blip-down images) may help minimize the impact of artifacts (e.g., (Gallichan et al., 2010; Mohammadi et al., 2012)).

Diffusion coefficients are measured by solving the signal decay equation for ADC using a reference image (typically b=0 s/mm^2^) and a diffusion-weighted image (typically b=~1000 s/mm^2^). Higher b-values increase diffusion sensitivity, and are recommended for HARDI methods, but they reduce signal available for measurement. Therefore, high b-value acquisitions require longer scan times or reduced image resolution to boost SNR and compensate for the lost signal. Multiple non-zero b-values are recommended for advanced methods like DKI or NODDI, which will further increase scan time compared to DTI. Early DTI neurodevelopment studies acquired only 6 directions (Evans, 2006; Lebel et al., 2008), whereas most recent DTI neurodevelopment studies acquire data with 30 or more diffusion encoding directions. More directions generally provide more robust parameter estimates (Jones, 2004), and are necessary for advanced diffusion models, but 6-direction data can produce robust parameter estimates with sufficient averaging (Lebel et al., 2012a).

Fine spatial resolution is desirable for measuring smaller structures and mitigating partial volume effects, which occur when multiple tissues within one voxel are averaged together to produce erroneous parameter estimates (e.g., WM is averaged with surrounding GM and CSF to produce artificially low FA and high diffusivity values) (Alexander et al., 2001; Vos et al., 2011). However, finer spatial resolution reduces the signal available for measurement in a voxel, and thus must be balanced by altering other parameters, such as increasing averages (i.e., increased scan time), or reducing brain coverage. In addition, inference from CSF signal can be suppressed via e.g. Fluid Attenuated Inversion Recovery-DTI, which improve tractography and quantification of diffusion parameters (Concha et al., 2005), but also come at a cost of increased scan time. Isotropic resolution is now common in DTI studies as it offers advantages for tractography, though interpolation and/or subsampling strategies can help compensate when the resolution is not identical in all dimensions.

Hardware is not easily modified at a given site, but will necessarily impact image quality and limit pulse sequence design. For example, higher field strength MRI scanners provide more signal for measurement, thus providing more flexibility for shorter acquisitions, finer spatial resolution, and higher diffusion weighting. Stronger gradients also permit higher b-values in shorter echo times, which provides more signal for measurement. Radiofrequency (RF) coils with multiple receive channels provide higher SNR than volume coils and allow parallel imaging, which reduces acquisition times and reduces distortions (e.g., eddy currents).

Children, particularly young children, can be anxious around the MRI scanner and have difficulty remaining still for long periods of time. dMRI is sensitive to motion, and participant cooperation is necessary to ensure high quality images that can be accurately and robustly measured. A good quality DTI sequence can be acquired in ~5-8 minutes, typical of the studies reviewed here, and is generally well-tolerated by older children and adolescents. Studies in younger children may benefit from even shorter acquisition sequences to improve subject compliance. The first step to reducing potential data confounds due to head motion is to improve each participant’s compliance with scanning procedures. Unfortunately, most developmental and clinical studies using dMRI fail to include, or report, rigorous training methodology use to actively reduce in-scanner head motion. Accepted procedures include the use of a mock scanner, behavioral training, and prospective motion correction (Hallowell et al., 2008; Theys et al., 2014). Mock scanning is the most common approach used to familiarize children and adolescents with the MRI environment (de Bie et al., 2012; de Bie et al., 2010; Hallowell et al., 2008; Raschle et al., 2011). Unfortunately, this often requires children to visit the hospital or laboratory on several occasions or increase the duration of the study visit. Recent training protocols have been described (Theys et al., 2014) that make the MRI session a pleasant experience for children, are time and resource efficient, do not require multiple visits, and most importantly, may improve the success rate of dMRI. To further improve compliance, accelerated acquisition sequences can be used to minimize time in the scanner, and/or prospective motion correction techniques can be used to compensate for motion during the scan, where available (Aksoy et al., 2011; Alhamud et al., 2012; Benner et al., 2011).

### 2.3. Diffusion image analysis

One of the simplest methods for quantifying diffusion parameters across the brain is region-of-interest (ROI) analysis. ROIs are drawn manually or automatically on images to isolate specific brain areas, and diffusion parameters are averaged across an ROI to provide one value per subject per region. ROI methods can be especially useful for measuring cortical or subcortical GM parameters, which otherwise have FA values that are too low for tract-based methods. Manual ROI analysis is user-dependent, especially when using small ROIs, though several studies report very high reliability (ICC>0.8) of FA, MD, AD, and RD (Bonekamp et al., 2007; Pfefferbaum et al., 2003). To ensure consistency across a study, all analysis should be done by the same operator, following strict criteria with respect to placement in each subject. This makes ROI analysis incredibly time-consuming, and unfeasible for large studies. Automated ROI methods register subject data to an atlas (e.g., JHU ICBM DTI-81 atlas), where brain regions can be queried automatically using the atlas labels. While automated methods overcome the laboriousness and user-dependency of manual ROI analysis, they rely on normalization to obtain diffusion parameters. Even small misregistration errors can bias results, in some regions more than others (Klein et al., 2009; Snook et al., 2007), and the adult templates that are typically used (pediatric DTI templates are not readily available) may not be appropriate for pediatric studies (Wilke et al., 2002; Yoon et al., 2009).

Voxel-based analysis (VBA) can be used to query diffusion parameters across the whole brain without *a priori* hypotheses about specific structures. This approach is automated, and thus user-independent, although it does depend on parameter choices. Generally, voxel based approaches normalize individual data to a template (standard or study-specific), smooth data or project it onto a skeleton, and then query individual voxels across all subjects to look for group differences or correlations. As with automated ROI methods, normalization can be problematic for VBA, especially in particular brain regions. Small registration errors can be mitigated using several different strategies. Smoothing blurs the edges of structures and boosts SNR, though smoothing can be problematic for heterogeneous DTI data, and results depend heavily on parameter choice (Jones et al., 2005). Thresholds (e.g., FA>0.25) can also be used in VBA to eliminate voxels that do not primarily contain WM, helping to ensure that the same tissue class is being compared across subjects. Tract-based spatial statistics (TBSS) (Smith et al., 2006) is a common voxel-based method, which creates a WM skeleton through the center of the WM and projects the highest FA values in the vicinity onto the skeleton. This helps overcome misregistration, but also eliminates much of the WM from analysis. TBSS is not necessarily tract-specific (i.e., projected FA values may be from different tracts) (Bach et al., 2014), and suffers from statistical bias along the skeleton (Edden & Jones, 2010). Because each voxel is analyzed separately, there are thousands or tens of thousands of statistical tests made during VBA. Thus, multiple comparison correction is essential to control for false positives, and most analysis programs have built-in multiple comparison correction tools. However, multiple comparison correction comes with a cost of false negatives. Thresholding and skeletonization also help control false positives by analyzing only a portion of the voxels in the brain, and thus reducing the number of comparisons, but they also reduce the brain areas queried. A few studies consistently find very high reliability of FA, MD, AD, and RD for VBA/TBSS using the same preprocessing pipeline (Madhyastha et al., 2014; Vollmar et al., 2010), but other reports suggest that VBA results are quite sensitive to the specific choices made for imaging smoothing and image coregistration, which can affect generalizability of the results (Jones et al., 2013). A recent reliability study of TBSS indicates that the across-session test-retest reproducibility errors are largely consistent across many different acquisition sites/vendors and within the range of 2–6% for all diffusion metrics (Jovicich et al., 2014), despite substantial differences across MRI vendors. In addition, this report found the most reproducible DTI metrics were FA and AD, followed by MD, and finally RD. Specific image analysis procedures, particularly the use of a common template and median filter smoothing, have been shown to markedly improve the reliability of TBSS (Madhyastha et al., 2014).

Tractography is another approach to the analysis of dMRI data (**Figure 1**). Tractography virtually reconstructs WM pathways, forming a three dimensional volume of interest for analysis. Deterministic tractography follows the primary diffusion vector from voxel to voxel using an FA threshold to remain in the WM and an angle threshold to avoid unlikely turns (Basser et al., 2000; Mori et al., 1999). Deterministic tractography can be prone to errors in regions of crossing fibers, where the tensor only models one fiber direction per voxel and FA values become artificially low. Probabilistic tractography takes into account the uncertainty of fiber directions in each voxel, providing a range of possible pathways and their likelihood (Behrens et al., 2003). Constrained spherical deconvolution (CSD) uses HARDI data to estimate multiple fiber populations within each voxel, which can then be used for tractography (Tournier et al., 2007). While CSD provides better fiber reconstructions than DTI (Farquharson et al., 2013), image acquisition and analysis requires significantly more time and processing capacity. Regardless of the method used, tractography provides a volume of interest from which to measure parameters that (presumably) reflect WM structure. Diffusion parameters can then be averaged across the entire WM tract to provide one value, which assumes some homogeneity across the tract, or evaluated over smaller portions of the tracts (Chen et al., 2016; Colby et al., 2012; Yeatman et al., 2014). Tractography loses some of the sensitivity to local changes if parameters are averaged across a whole tract, but introduces multiple comparison problems if too many smaller sections are examined individually.

**Figure 1.**
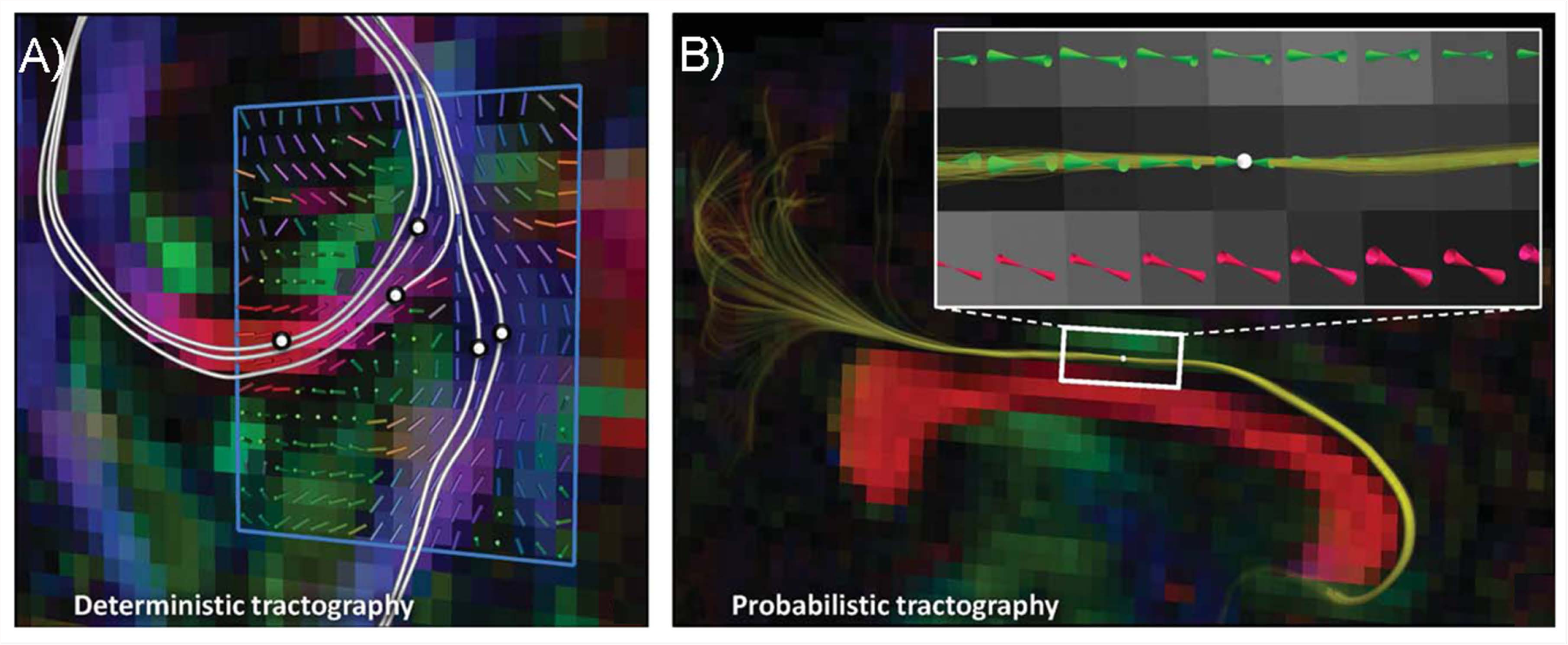
Conceptual examples of deterministic and probabilistic tractography based on the diffusion tensor model. A) Example of deterministic tractography where the white lines represent fiber tract pathways that were reconstructed by following the principal diffusion directions in consecutive steps, initiated bidirectionally at the indicated locations (seed points). For each of the pathways, there is no information available about the precision/dispersion that is associated with their tract propagation. B) Example of probabilistic tractography where the set of multiple lines provides a feel for the degree of uncertainty related to the tract reconstruction initiated from the single seed point. Adapted with permission from Tournier et al. (2011).

As with ROI analysis, tractography methods can be automated to reduce the laboriousness of manually tracking fibers in a large population (O’Donnell & Westin, 2007; Zhang et al., 2010). Semi-automated methods where a study-specific set of seeding and target regions are created and used to automate tracking have been used to study neurodevelopment (Lebel et al., 2008), and as automated tractography methods become more widely available (e.g., FreeSurfer’s TRACULA, (Yendiki et al., 2011; Yendiki et al., 2016)), they are likely to be used in future neurodevelopment studies. Automated tractography methods can be more forgiving of registration errors and inter-subject variability than atlas-based ROIs or voxel-based analyses. This is because multiple seeding and target regions can be used that are each larger than the tract of interest, but together combine to select only fibers from the desired pathway.

Finally, a completely different approach to diffusion data is to create networks and conduct a graph theory analysis. Graph theory considers the brain as a whole network, defining nodes and edges to create a model of the brain’s structural connectivity, and then examining its topological and geometrical properties (Bullmore & Bassett, 2011). Quantitative measures of network connectivity include degree (the number of edges emanating from a node), path length (the length of each edge), and clustering coefficient (the efficiency of local connectivity). Brain networks exhibit small-world properties, meaning that the clustering coefficient is greater than that of a random network (Bullmore & Bassett, 2011). Small-worldness and other measures such as modularity or efficiency can be quantified and examined with respect to age to measure development. A variety of software packages are available to conduct graph theory analysis of brain networks (e.g., the Brain Connectivity Toolbox, (Rubinov & Sporns, 2010)). In dMRI, nodes are often defined as cortical or subcortical brain regions, and the edges are defined as the WM pathways that connect these nodes. Tractography is typically used to determine the presence or absence of an edge between two nodes defining a binary network. Analysis can then proceed on this unweighted network, or edges can be weighted by a measure of their connectivity (e.g., FA, 1/MD, number of streamlines) for further analysis. Graph theory is powerful for analyzing large volumes of data, and can provide unique insight into developmental changes. However, there are multiple ways of defining a network from a diffusion weighted image, and findings using e.g., FA weighted networks can be very different from findings on weighted graphs using streamline count. Furthermore, network parameters may be influenced even by a small number of false positive connections (which are common), and measures are sensitive to parameter choices (e.g., edge weighting, connection thresholds) (Drakesmith et al., 2015).

All analysis approaches for dMRI data have distinct advantages and disadvantages. Thus, interpretation of findings must be made with caution and a full understanding of how both image acquisition and analysis choices may influence results. Ultimately, it is only through replication across studies with different populations, image acquisition, and analysis strategies that we can piece together the full picture of brain development during childhood and adolescence.

### 2.4. Quality control and motion

Once high quality dMRI data is collected, practical challenges remain that affect the reliability and reproducibility of the results (Le Bihan et al., 2006). The quality of diffusion measurements is susceptible to eddy currents, echo-planar distortions, rotation errors in the b-matrix, partial volume effects, scanner artifacts (e.g., noise spikes) and susceptibility artifacts (Anderson, 2001; Bastin et al., 1998; Pasternak et al., 2009; Skare et al., 2000b). But, perhaps the largest data confound in human dMRI samples is head motion (Liu et al., 2015; Power et al., 2012), especially in studies of children (Yoshida et al., 2013) and adolescents (Roalf et al., 2016; Satterthwaite et al., 2012; Yendiki et al., 2014). For example, head-motion was found to induce group differences in DTI measures between children with autism and typically developing children (Yendiki et al., 2014) and attenuate the relationships between age and FA and MD in a developmental sample (Roalf et al., 2016) (**Figure 2**). These confounds likely contribute to inaccuracies in the tensor fitting of diffusion data (Le Bihan et al., 2006), have come under scrutiny (Heim et al., 2004; Jones et al., 2013; Jones et al., 1999; Lauzon et al., 2013; Leemans & Jones, 2009; Owens et al., 2012; Tournier et al., 2011; Vos et al., 2011; Yendiki et al., 2014), and are subsequently the focus of several new methods seeking to mitigate their impact (Li et al., 2014; Li et al., 2013; Oguz et al., 2014).

**Figure 2.**
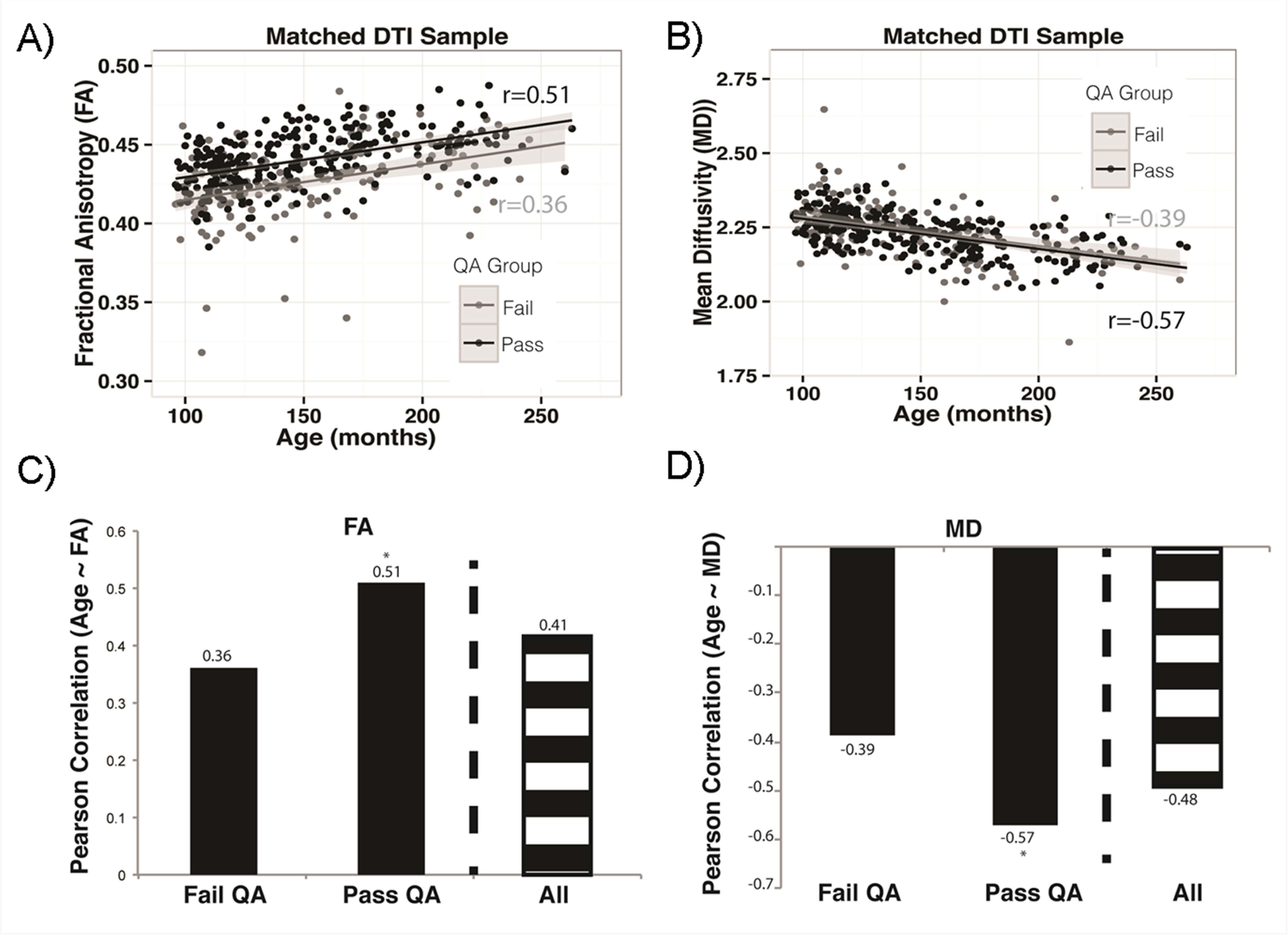
Age-related differences in diffusion tensor imaging (DTI) metrics in individuals aged 8-21 years in data that fails or passes quality assessment (QA). A) Data that failed QA had a significantly lower correlation between age and FA than data that passed QA. B) Data that failed QA also had a significantly lower correlation between age and MD than data that passed QA. C) & D). Summary plots of the correlations between age and DTI metrics in data failing QA, passing QA and all data combined. Adapted with permission from Roalf et al. (2016).

Surprisingly, most studies do not report procedures for diffusion data quality control (QC) and its impact on diffusion metrics. Typically, diffusion studies report that data with “obvious artifacts” are removed or excluded. However, a transparent, standardized estimate of quality assurance is rarely given. It is likely that unaccounted artifacts result in suboptimal tensor estimation and may negatively influence commonly derived diffusion scalar metrics and estimates of tractography. Few clinical or neurobiological studies report specific artifacts or quantify SNR or motion with diffusion findings despite the availability of prior methodological studies which have outlined the influence of problematic diffusion data on typical outcome metrics (Anderson, 2001; Armitage & Bastin, 2001; Bastin et al., 1998; Chen et al., 2015; Heim et al., 2004; Jones, 2004; Jones & Basser, 2004; Pierpaoli & Basser, 1996). These studies demonstrate that systematic data loss or the exclusion of data from too many diffusion directions can affect the estimates of FA, and to a lesser degree MD (Chen et al., 2015; Jones, 2004; Jones et al., 1999). These methodological studies tend to be small and use simulated data, but confirm that artifacts in diffusion data can be overcome if enough directions are collected, or if the loss of data during a given diffusion acquisition is random, especially across subjects (Chen et al., 2015; Heim et al., 2004; Jones, 2004; Jones & Basser, 2004). In addition, because data quality is often systematically related to a phenotype of interest (e.g., age, diagnosis, cognition, symptom severity) and data quality is inherently subject dependent (e.g., correlation between age and motion), low quality data has the potential to obscure the presence of real effects or produce spurious associations with study phenotypes.

Despite the dangers of poor quality data on result validity, automated measures for QC of diffusion data remain limited. Manual inspection of multivolume diffusion data is time consuming, subject to operator bias, and translates poorly to large-scale imaging studies. Studies of noise in diffusion provide a useful framework for identifying how such noise affects diffusion properties (Ding et al., 2005; Farrell et al., 2007; Hasan, 2007; Skare et al., 2000a). Several recent studies indicate promise for implementing automatically derived quality assurance metrics that reduce the amount of manual QC effort, including measures of SNR and the use of outlier detection, to quantify data quality prior to image processing (Lauzon et al., 2013; Li et al., 2014; Li et al., 2013; Oguz et al., 2014). However, much of this work has used relatively small samples or simulated data, and none have focused primarily on an adolescent neurodevelopment sample (although (Lauzon et al., 2013) present data in a large pediatric sample). Recent work in a developmental sample showed, not surprisingly, that low-quality DTI data had significantly lower FA throughout major WM tracts and showed higher MD in several brain regions as compared to data that passed rigorous quality assurance (Roalf et al., 2016). Manual visualization indicated that over 10% of the data had significant artifact, which was confirmed by automatic QC detection algorithms. While high, this is a smaller percentage than in some pediatric samples (Li et al., 2013), and is likely representative of what could be expected in adolescent samples. Automated tools to investigate and correct DTI artifacts, such as DTIPrep and RESTORE (e.g., (Chang et al., 2005; He et al., 2014; Li et al., 2014; Liu et al., 2015; Oguz et al., 2014), aim to investigate, eliminate and/or correct problematic slices or volumes in diffusion data. Other tools exist to quickly and automatically interrogate diffusion data quality (Roalf et al., 2016).

### 2.5. Longitudinal image analysis: promise and challenges

Longitudinal data is especially critical for understanding brain development, as it can demonstrate the within-subject changes that cross-sectional data can only suggest. Longitudinal analyses have increased power compared to cross-sectional studies, as inter-subject variability is reduced in favor of intra-subject measurements of changes over time. However, longitudinal studies can be challenging, and the utility of dMRI to measure longitudinal change in the developing brain requires reliability and stability of measurement. Not surprisingly, variation in acquisition, preprocessing, or analysis can all affect reliability of the diffusion signal, and subsequently its interpretation (Madhyastha et al., 2014). Ideally, longitudinal diffusion studies would occur using the same scanner, head-coil, and protocol. While achieving consistency in all of these factors is possible over a short to medium time frame (for example see (Roalf et al., 2016)), it is all but inevitable that systematic changes or upgrades will occur causing one or many of these factors to change over the long term. Unfortunately, changes in the MRI system or protocol may affect longitudinal interpretation of the data. That is, plausible biological differences in diffusion measurements may in fact be related to technical factors associated with hardware platforms (scanner vendor), software releases, and specific imaging parameters, including different pulse sequences, parallel imaging techniques, and reconstruction algorithms associated with a specific vendor (Wang et al., 2016). In addition, other acquisition parameters may differ substantially including, receiver bandwidth, echo time (TE), and slice thickness. Changes in the RF head coils can differentially affect SNR, and since diffusion imaging is intrinsically a low SNR method (Skare et al., 2000b), it is imperative to optimize the experimental conditions and maintain these conditions as best as possible. Differences between MRI scanner vendors are expected, however, scanners of the same model and software release can also yield different results (Vollmar et al., 2010). Thus, in general, ad-hoc mixing of diffusion data collected from different scanners or acquired using different protocols should be avoided.

Given the duration needed for longitudinal studies to elucidate developmental changes, it is often not possible to achieve consistency across all of the factors noted above. One powerful approach to conduct comprehensive studies of basic neuroanatomy and measure developmental change is through the use of multi-center studies (Lemkaddem et al., 2012). This approach employs common data collection and analysis strategies, distributes the data acquisition load across multiple sites, substantially speeds up the research process, and increases participant access to the study, which subsequently increases generalizability of the findings. Some examples of this approach include Alzheimer’s Disease Neuroimaging Initiative (ADNI (Van Horn & Toga, 2009)), the HCP (Van Essen et al., 2012), and the recently started Adolescent Brain Cognitive Development (ABCD) study. This approach embraces the known variability in scanner platform, while attempting to systematically control as many confounds as possible, and provides an opportunity to move toward standardized diffusion protocols that are optimal across platforms and protocols. It requires uniform QC procedures to ensure that data from one site or scanner does not create bias in analysis, exclusion of substandard data, and detailed methods to maintain compliance (e.g. phantom studies).

As noted above, combining data from different scanner vendors will introduce additional sensitivity to systematic inter-site variability. However, there is accumulating evidence (Cercignani et al., 2003; Danielian et al., 2010; Magnotta et al., 2012; Pagani et al., 2010; Pfefferbaum et al., 2003; Vollmar et al., 2010) that inter-site variability can be low (~5%), particularly in large WM tracts (Fox et al., 2012; Grech-Sollars et al., 2015; Magnotta et al., 2012; Teipel et al., 2011; Wang et al., 2016; Zhu et al., 2011). Depending upon vendor and methodology, the coefficient of variation for both within-site and across-site diffusion metrics be as low as 0.5% and 2%, respectively (Magnotta et al., 2012). This is particularly encouraging for longitudinal studies that occur using the same scanner, but also suggests that subtle changes in WM structure can be measured longitudinally when changes occur to the MRI scanner or protocol, given certain limitations.

Reliability of diffusion metrics is paramount for detecting within-person change in longitudinal studies. Fortunately, the estimated reliability of diffusion metrics taken within one year is quite high (~90%), but decreases slightly (~80%) over longer intervals (Shou et al., 2013; Zipunnikov et al., 2014). Thus, diffusion metrics appear to be quite stable, especially when compared to fMRI (Shou et al., 2013), although they are generally more variable than T1-weighted structural MRI measures (e.g., volume, cortical thickness) (Morey et al., 2010; Shou et al., 2013). Further, both reliability and sensitivity have been shown to improve by longitudinal pipelines reconstructing WM pathways jointly using the data from all available time points (Yendiki et al., 2016). Unfortunately, specifically developed longitudinal processing pipelines are not yet available for several of the most popular software packages (but see (Aarnink et al., 2014; Yendiki et al., 2016)).

Another challenge in longitudinal studies is statistical data analysis. Longitudinal statistics are more complex than cross-sectional analyses, and most available standard analysis programs are not set up for easy analysis of longitudinal data. One approach for longitudinal data analysis is to use paired t-tests or repeated measures analyses where first and subsequent scans are compared to each other within individuals. This is the most statistically powerful approach, but it does not provide much information about how rates of change vary across ages or brain regions. Brain development is nonlinear, so paired t-tests and repeated measures are not ideal for studies with wide age ranges. They are also not appropriate if the time interval between scans varies greatly, or if there are different numbers of scans for each participant. In these cases, a mixed-models analysis approach may be more appropriate. This is a general linear model, where subject is included as a random factor, alongside covariates of interest such as age and sex. In this way, a trajectory of age-related changes can be calculated while also considering the shared variance within each subject. For more in-depth introductions and discussions of issues related to longitudinal modelling, we refer to other papers in this special issue.

Despite challenges in acquisition and analysis, the advantages of longitudinal analysis far outweigh its challenges. Longitudinal studies have, as reviewed below, provided great insight into development during childhood and adolescence, and have (unsurprisingly) proven themselves to be more sensitive to age-related changes than cross-sectional studies. Future longitudinal studies following children over longer periods of time and/or with more data points have great potential to elucidate the developmental trajectories of WM within individuals and relate these to cognition, affect and behavior.

## 3. Diffusion MRI: Applications to understanding brain development in childhood and adolescence

### 3.1. Developmental changes in dMRI parameters

The major fiber pathways in the brain are already present and identifiable at birth, but very rapid changes in DTI indices of WM microstructure are seen across infancy (for reviews see (Dubois et al., 2014; Qiu et al., 2015)). For instance, a large study including 211 young children and 295 scans indicated that in the first two years of life, FA in ten major WM tracts increases by 16-55%, RD decreases by 24-46%, and AD decreases by 13-28%, with faster changes in the first year than in the second for all tracts investigated (Geng et al., 2012). Such massive changes are not surprising given the enormous behavioral and psychological development seen in this period of life. Studies focusing on early life brain development will however not be covered in the present review, as the nature and scale of these changes are very different from those seen in later childhood and adolescence. In addition to the methodological challenges for dMRI studies of children and adolescents discussed in this review, studies of infants and young children have other specific major challenges, including particular difficulties in obtaining relatively motion-free images from babies (as sedation is not used without clinical indication), image registration and alignment, use of adult-based brain atlases, and dramatically changing image intensity contrasts which may lead to misclassification of tissues, especially when using automated software, and make comparisons across age difficult (Sled & Nossin-Manor, 2013).

Numerous cross-sectional studies have been carried out to investigate age-related differences in DTI parameters in children and adolescents, which, despite sample and methodology differences, consistently demonstrate FA increases and overall diffusivity decreases with increasing age in most WM regions (e.g., (Asato et al., 2010; Ashtari et al., 2007; Barnea-Goraly et al., 2005; Ben Bashat et al., 2005; Clayden et al., 2012; Eluvathingal et al., 2007; Giorgio et al., 2008; Klingberg et al., 1999; Lebel et al., 2010; Lebel et al., 2008; Muftuler et al., 2012; Pohl et al., 2016; Qiu et al., 2008; Schmithorst et al., 2002; Snook et al., 2005; Tamnes et al., 2010; Wu et al., 2014; Yu et al., 2014), for reviews see (Cascio et al., 2007; Schmithorst & Yuan, 2010), for a meta-analysis see: (Peters et al., 2012)).

Longitudinal DTI studies of children and adolescents are becoming more common, but are still scarce (**Table 1**). In contrast to the cross-sectional studies, longitudinal studies track changes over time within individuals and can directly relate these estimates to influencing factors, outcomes or concurrent developmental changes for instance in cognition, social and affective processing or symptomatology. Most of the available studies have employed accelerated longitudinal designs which allow for investigation of wide age-ranges over shorter duration data collection periods, but with the trade-off being the inherent missing data since each subject’s measurement schedule covers only part of the age-range of interest (Galbraith et al., 2017). One study employing a single cohort longitudinal design has also been performed (Brouwer et al., 2012); focusing on a narrow age range, but alleviating the missing data issue since each individual is followed over approximately the same period. Notably, all available longitudinal studies include only two observations for all or the majority of the participants, and this remains a major limitation of the field, as such datasets do not allow for optimal modelling of non-linear within-person change.

**Table 1.**
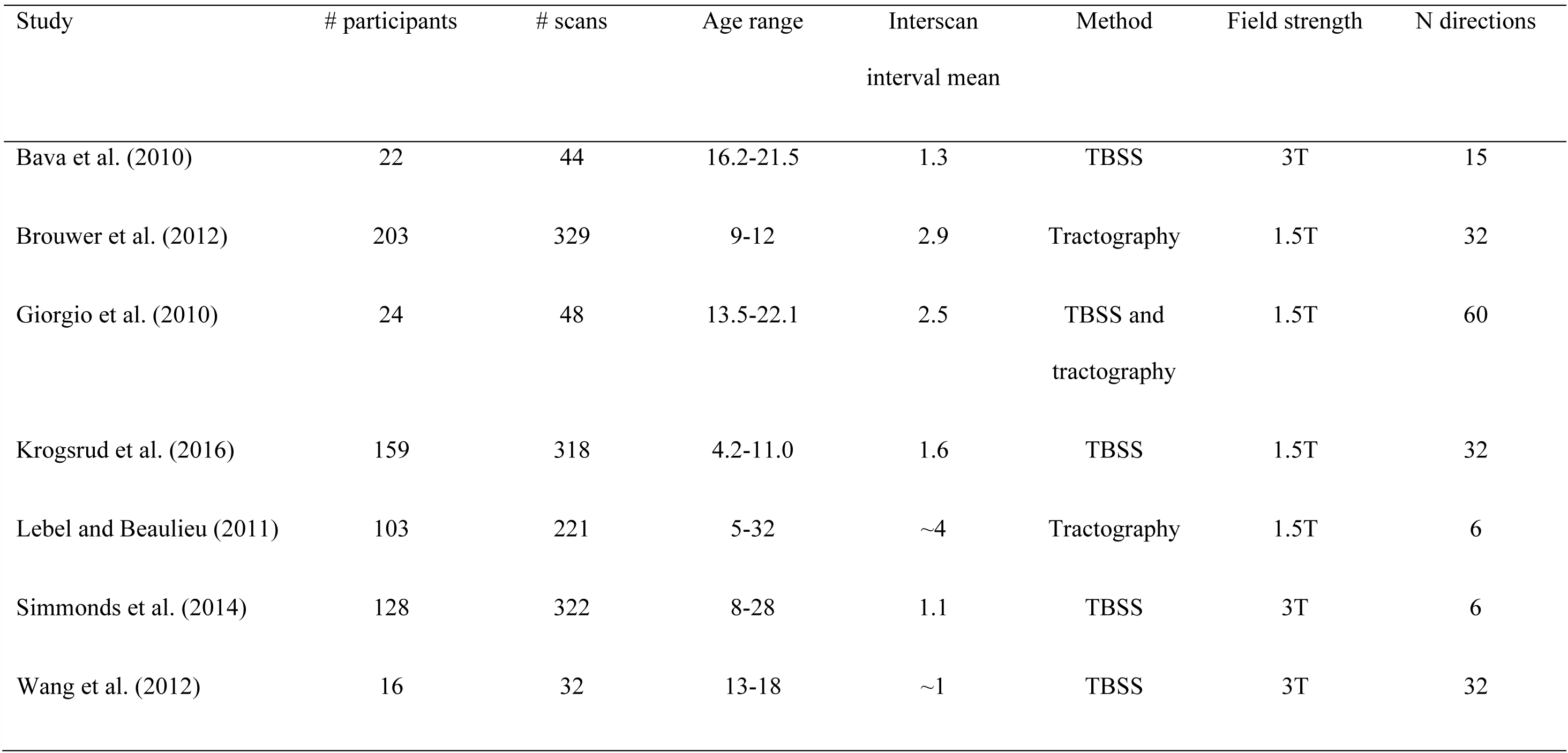
Longitudinal DTI studies of white matter microstructure development in childhood and adolescence

| Study | # participants | # scans | Age range | Interscan<br>interval mean | Method | Field strength | N directions |
| --- | --- | --- | --- | --- | --- | --- | --- |
| Bava et al. (2010) | 22 | 44 | 16.2-21.5 | 1.3 | TBSS | 3T | 15 |
| Brouwer et al. (2012) | 203 | 329 | 9-12 | 2.9 | Tractography | 1.5T | 32 |
| Giorgio et al. (2010) | 24 | 48 | 13.5-22.1 | 2.5 | TBSS and<br>tractography | 1.5T | 60 |
| Krogsrud et al. (2016) | 159 | 318 | 4.2-11.0 | 1.6 | TBSS | 1.5T | 32 |
| Lebel and Beaulieu (2011) | 103 | 221 | 5-32 | ~4 | Tractography | 1.5T | 6 |
| Simmonds et al. (2014) | 128 | 322 | 8-28 | 1.1 | TBSS | 3T | 6 |
| Wang et al. (2012) | 16 | 32 | 13-18 | ~1 | TBSS | 3T | 32 |

The first published longitudinal DTI studies were of very small scale with limited power to detect true effects, and more vulnerable to effects of outliers. Giorgio et al. (2010) analyzed data from 24 adolescents in the age-range 13-22 years and used TBSS to obtain DTI measures from the WM skeleton of scans collected at two time-points on average 2.5 years apart, as well as probabilistic tractography to isolate selected tracts. Their results showed bilateral significant FA increases in widespread regions in the WM skeleton and the arcuate fasciculi, but not the corticospinal tracts. The FA increases were mainly driven by increases in AD, while RD remained relatively unchanged. The same year, Bava et al. (2010) published results based on TBSS analyses of a dataset consisting of two time-points with a shorter interval (mean 1.3 years) from 22 slightly older adolescents (age-range 16-21 years). The results showed significant FA increases over time, but only in limited regions in the right hemisphere. Additionally, and in contrast to the findings by Giorgio et al. (2010), Bava et al. (2010) also observed decreases in both RD and AD in multiple regions.

A larger study published by Lebel and Beaulieu (2011), included 103 participants in a much broader age-range, 5-32 years, and with 2-4 scans per participant (mean = 2.1) acquired over 1-6 year intervals. Deterministic tractography for the following 10 major WM tracts was performed: genu, body and splenium of the corpus callosum, corticospinal tract, superior and inferior longitudinal fasciculi, superior and inferior fronto-occipital fasciculi, uncinate fasciculus, and cingulum. All showed significant nonlinear (quadratic) developmental trajectories, with decelerating increases for FA values (**Figure 3**) and decelerating decreases for MD, except the corticospinal tract which had linearly decreasing MD. These changes were due primarily to decreasing RD. Another large tractography study, but with a different type of design and sample, was published by Brouwer et al. (2012). This was a single cohort longitudinal twin study including 203 individuals with 1-2 scans per participant (mean = 1.6), with the baseline scans acquired at age 9 and the follow-up scans at age 12. FA increased in all fiber tracts investigated, with average annual change rates of 0.4-2.3%. Most tracts showed increases in AD and decreases in RD, with change rate in RD being greater than that of AD in 9 of the 14 tracts that were studied, and in the overall WM mask. So far, only one longitudinal study with more than two time-points for many participants has been published. Here, Simmonds et al. (2014) followed 128 participants with 1-5 scans (322 scans in total, mean scans = 2.5, 60/128 participants with > 2 scans) acquired approximately annually in the age-range 8-28 years. Across the WM skeleton and across atlas-defined ROIs, increases in FA and decreases in RD with age were seen. Across the skeleton, AD also decreased with age, but AD did not significantly change with development in the majority of ROIs.

**Figure 3.**
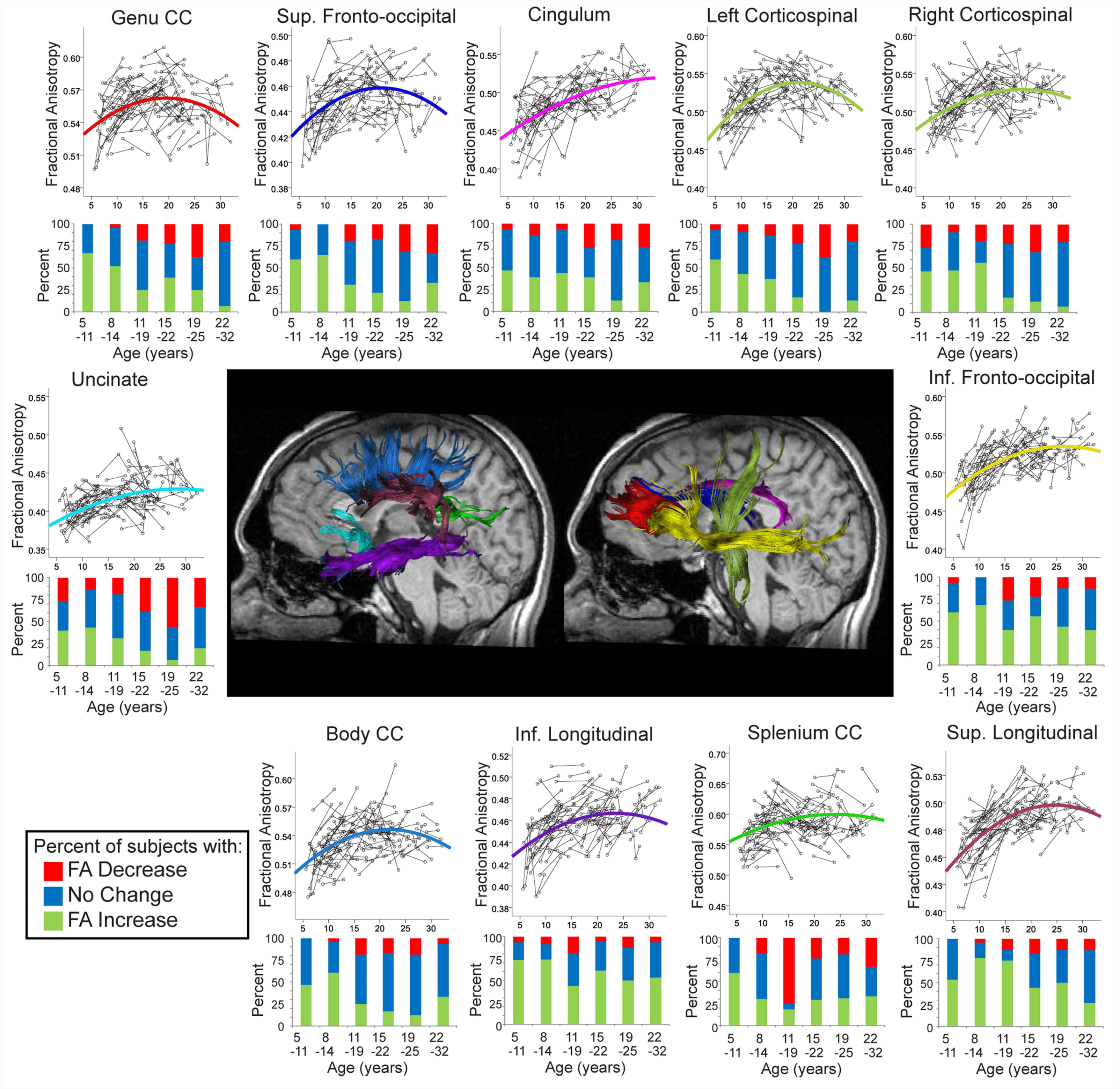
Longitudinal age-related changes of fractional anisotropy (FA) in healthy individuals aged 5- 32 years. Spaghetti plots with the best fitting models and bar graphs depicting the percentage of subjects whose FA increased (green), decreased (red), or did not change (blue) in six age groupings are shown for different WM fibers, derived using a deterministic tractography method. Adapted with permission from Lebel and Beaulieu (2011).

In a recent unique study focusing on the preschool and early school years, specifically ages 4-11 years, Krogsrud et al. (2016) analyzed longitudinal DTI data from 159 participants scanned twice. Across the averaged WM skeleton, FA showed a significant linear increase over time, while there were linear decreases in MD and RD, and AD showed only a weak decrease. Some regional differences were observed, although the same general pattern was seen for most of the investigated atlas-defined tracts. These mostly linear WM microstructural changes during middle and late childhood complement the above described findings by Lebel and Beaulieu (2011) of nonlinear trajectories across adolescence.

In summary, the few available longitudinal DTI studies generally confirm the findings from the cross-sectional studies, documenting continued WM microstructural development though childhood and adolescence with decelerating bilateral and widespread increases in FA and decreases in MD and RD, while the conclusion for AD is less clear. It is not possible, either on a group or individual level, to specify an age at which the brain becomes ‘fully developed’. DTI indices of WM microstructure are never static, but rather reflect lifelong maturation, experience-dependent plasticity and degeneration. Although a certain degree of stability can be seen in adulthood relative to childhood and adolescence, there are no periods of life in which brain WM microstructure is fixed. DTI studies including participants across major parts of the human lifespan have extended the findings from the developmental studies, indicating non-monotonic age trajectories of FA, MD and RD characterized by three phases: (1) initially fast, but decelerating changes through childhood and adolescence and into early adulthood followed by (2) relative stability in mid-adulthood, with subsequent (3) accelerating changes in senescence (Lebel et al., 2012b; Sexton et al., 2014; Westlye et al., 2010).

Notably, DTI can also be used to investigate developmental changes in tissue microproperties in subcortical GM structures and the cerebral cortex. Cross-sectional results from a large sample of children and adolescents by Lebel et al. (2008) for instance indicate greater magnitudes of age-related DTI changes in deep GM structures, specifically for FA increases in regions including the caudate nucleus, globus pallidus and putamen, than in WM tracts (see also (Pal et al., 2011)). Grydeland et al. (2013) investigated intracortical MD in a large cross-sectional lifespan sample and observed linear decreases in large frontal and temporal regions in childhood and adolescence. However, DTI measurement in cortical and subcortical GM is challenging, particularly for thin cortical and small subcortical regions, due to the relatively low resolution of standard DTI sequences, which likely result in partial-volume effects (Grydeland et al., 2013; Koo et al., 2009).

Developmental studies using more recent and advanced dMRI models such as DKI and NODDI are also becoming more common, yet only cross-sectional studies are so far available. Several DKI studies of children and adolescents have shown that mean kurtosis shows age-related increases (Das et al., in press; Falangola et al., 2008; Grinberg et al., 2017). A rare developmental NODDI study by Chang et al. (2015) including 66 participants 7-63 years old indicate that the age-related FA increase during childhood and adolescence is dominated by increasing NDI, which points specifically to increases in fiber diameter and myelination, while the decrease in FA later in life may be driven by increasing ODI (**Figure 4**). Moreover, the results from this study indicated that NODDI metrics predicted chronological age better than DTI metrics, a conclusion also recently supported by Genc et al. (2017). Further work, including longitudinal studies, is needed to explore the potential of these promising techniques for increasing our understanding of brain development.

**Figure 4.**
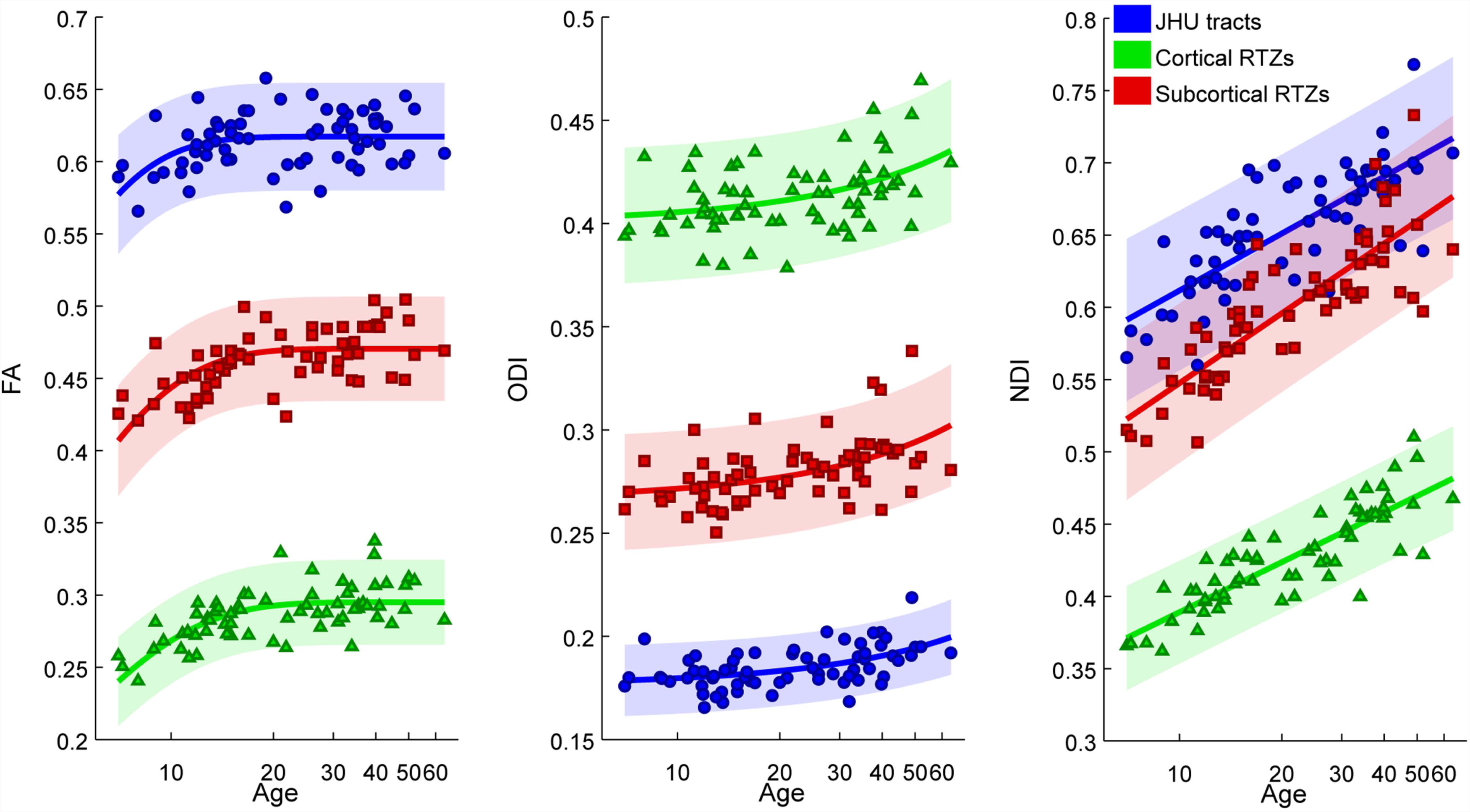
Age-related differences in average fractional anisotropy (FA), orientation dispersion index (ODI), and neurite density index (NDI) in the age-range 7-63 years across the WM skeleton in atlas defined tracts (JHU) (blue), cortical regional termination zones (RTZs) (green), and subcortical regional termination zones (RTZs) (red). Shaded regions represent 95% confidence intervals. Adapted with permission from Chang et al. (2015).

### 3.2. Sex differences and puberty

With sex-related differences apparent in total brain volume and volumetric analyses of grey and white matter (Mills et al., 2016; Ruigrok et al., 2014), it is important to consider whether, and how, WM microstructure may differ between the sexes, and each of the longitudinal studies mentioned above have sought to address this question with variable findings. Giorgio et al. (2010) found no sex differences in FA across the WM skeleton, noting that their study was not adequately powered. Contrastingly, Wang et al. (2012) reported higher global diffusivity values (MD, RD and AD) in females compared to males, and also an age-by-sex interaction on global mean FA, where FA was low in younger males, and significantly increased with age, while in females there was no change with age. Both these studies did however include small and likely underpowered samples. A larger study performed by Simmonds et al. (2014) found changes in WM microstructure in both sexes during adolescence, and there were age-by-sex interactions during both late childhood/early adolescence (8-12 years) and late adolescence (16-19 years), with males showing significantly larger FA increases than females. Lebel and Beaulieu (2011) reported regional differences in WM between the sexes, with females demonstrating higher FA in the splenium of corpus callosum, and higher MD in the cingulum, corticospinal tracts and superior fronto-occipital fasciculus compared to males, while males exhibited higher FA in the cingulum, bilateral corticospinal tracts, superior longitudinal fasciculus and uncinate fasciculus. While significant, the size of these sex differences was small (1-3%) and less than the variation seen between scans from the same individual (5-7%), as emphasized by the authors who advised interpreting these sex differences with caution.

One prominent hypothesis for the differential age trajectories between males and females of WM development during adolescence proposes that pubertal development, and the sex steroid hormones that drive puberty, may influence brain maturation, and differential hormonal exposure between the sexes may result in different growth patterns. A small number of cross-sectional studies have empirically studied this question (Asato et al., 2010; Herting et al., 2012; Menzies et al., 2015; Peper et al., 2015), and each supports a role for puberty in WM microstructural development, but the lack of longitudinal data and the range of WM indices analyzed prevent any clear consensus on the nature of this role.

### 3.3 Spatiotemporal patterns and network approaches

Importantly, the timing and rates of the DTI developmental changes vary regionally in the brain. From cross-sectional studies, a pattern in which major WM tracts with fronto-temporal connections develop more slowly than other tracts has emerged (Lebel et al., 2008; Tamnes et al., 2010). Further, cross-sectional studies with very wide age-ranges covering large parts of the life-span suggest that of the major fiber bundles, the cingulum which is implicated in cognitive control has a particularly prolonged development (Lebel et al., 2012b; Westlye et al., 2010). In an interesting cross-sectional study by Colby et al. (2011), broad regional gradients in the developmental timing of WM skeleton FA along both inferior-to-superior and posterior-to-anterior directions over the age-range 5 to 28 years were seen, indicating relatively late development of superior and anterior regions. Looking at longitudinal change rates along different slice-by-slice spatial gradients, Krogsrud et al. (2016) found support for a posterior-to-anterior gradient of change magnitude, with greater DTI changes seen in frontal regions during childhood. Analyzing within-subject changes within tracts, operationalized as an increase or decrease of greater than 1 SD as computed based on a separate interscan reliability test, Lebel and Beaulieu (2011) found that changes in DTI parameters were mostly complete by late adolescence for projection and commissural tracts, while post-adolescent development was indicated for both FA and MD in association tracts.

Beyond the characterization of microstructural tissue properties, DTI can, in conjunction with graph theoretical analysis, be used to investigate developmental changes in brain network organization. Although many features of complex networks like small-worldness, highly connected hubs (together forming a rich club) and modularity are already established at birth, they are thought to mature across childhood and adolescence at a macroscopic neuroimaging scale measured using for instance DTI-based weighted tract connections between multiple local GM regions (Vertes & Bullmore, 2015). Developmental dMRI studies have shown that global and local efficiency increases, clustering decreases, and modularity stabilizes or decreases with age, overall indicating that WM structural networks move from being local to more distributed and integrated (e.g., (Chen et al., 2013; Dennis et al., 2013; Koenis et al., 2015; Wierenga et al., 2016) for a recent systematic review see (Richmond et al., 2016)). Findings in a recent small longitudinal study also indicate a spatial refinement of connectivity between hubs during late adolescence (Baker et al., 2015).

### 3.4 Individual differences and atypical development

The majority of DTI studies have described developmental trajectories at a group level, overlooking the extensive differences in DTI measures between individuals at any given age, and the variability between individual developmental trajectories seen in longitudinal studies. This variation results from the continual interaction between genetic and epigenetic influences alongside external and environmental influences on the brain. Twin studies document substantial heritability for DTI indices of WM microstructure (Blokland et al., 2012; Brouwer et al., 2012; Kochunov et al., 2015) and overall structural networks (Koenis et al., 2015), but also that genetic influences vary with age (Chiang et al., 2011), and that changes in WM microstructure are also influenced by external factors.

Environmental and experiential variables influence WM microstructure across the lifespan. Prenatal factors including exposure to alcohol, cocaine, methamphetamines and tobacco can impact on early brain development with long-term consequences (Lebel et al., 2013; Liu et al., 2011; Roussotte et al., 2010; Treit et al., 2013; Warner et al., 2006). In more subtle ways, recent results also indicate that maternal mental health and emotional state have long-term neurobiological effects on the fetus (Lebel et al., 2016; Sandman et al., 2015). A meta-analysis investigating the impact of preterm birth on the WM microstructure from childhood to young adulthood identified a number of regions across the brain where differences in FA were seen (Li et al., 2015). Adversity in neonatal life and childhood can further impact on WM development with effects that persist into adolescence (Kumar et al., 2014; Schiller et al., 2017).

External influences during adolescence and young adulthood including sports-related concussion (Meier et al., 2016), use of alcohol, cannabis, tobacco and inhalants (Bava et al., 2013; Cookey et al., 2014; Gogliettino et al., 2016; Silveri et al., 2016; Yuncu et al., 2015) and sleep patterns (Telzer et al., 2015) have short and medium-term effects on WM development, although there is less evidence documenting whether these effects persist through adulthood. Multiple DTI studies document the effects of motor and cognitive training on WM microstructure, reporting FA increases and diffusivity decreases following interventions in both adults and children (Engvig et al., 2012; Gebauer et al., 2012; Krafft et al., 2014; Lovden et al., 2010). The durability of these effects, the potential of these changes to impact on other functional domains, and the effectiveness of these interventions in larger populations are still very much debated.

Studies assessing the impact of WM microstructural properties on cognitive, affective, motivational and social processes have demonstrated cross-sectional correlations across childhood and adolescence between WM development, particularly in fronto-parietal pathways, and a range of behaviors including response inhibition (Madsen et al., 2010), response time variability (Tamnes et al., 2012), working memory (Peters et al., 2014; Vestergaard et al., 2011; Østby et al., 2011), mnemonic control (Wendelken et al., 2015) and sustained attention (Klarborg et al., 2013). Cross-sectional studies like these assume linear age effects across childhood and adolescence, and a constant relationship between brain and behavioral indices. Few longitudinal studies have been performed, although a recent study by Achterberg and colleagues (2016) showed non-linear development of frontostriatal WM microstructural development between childhood and early adolescence, which predicted non-linear changes in delayed discounting ability over the same time period.

DTI and other dMRI models are also potentially powerful tools for detecting and charting abnormal WM microstructure and WM microstructure development in childhood and adolescence in mental health and neurological disorders, such as schizophrenia (Tamnes & Agartz, 2016), bipolar and unipolar depression (Serafini et al., 2014), ADHD (van Ewijk et al., 2012) and autism spectrum disorder (Aoki et al., 2013). However, the fact that lower anisotropy and higher diffusivity is observed in a wide range of different clinical groups (and brain regions) when compared to healthy controls, both in adults and in children, prompts us to question the specificity of such findings. Moreover, most available reports in pediatric populations are cross-sectional case-control studies – many with limited sample sizes – and often don’t include analyses of age-effects (but rather only matches or statistically controls for age).

Subclinical manifestations of most mental health disorders exist in the general population, including in children and adolescents (White, 2015), and neuroimaging population studies and studies of children and adolescents with subclinical symptoms are not only important for understanding risk-factors or early manifestations of disorders, but also appear to support a dimensional perspective on several brain phenotypes associated with mental health. For example, individuals showing subclinical symptoms for psychosis spectrum disorders (clinical high-risk) and individuals with diagnosed relatives (genetic high-risk) have brain microstructural changes at least somewhat overlapping with those reported in clinical groups (Arat et al., 2015; O’Hanlon et al., 2015; Peters & Karlsgodt, 2015), and a recent longitudinal study showed how higher levels of anxiety and depression symptoms was associated with regionally reduced rates of FA development in typically developing youth (Albaugh et al., 2016).

## 4. Future directions and conclusion

The longitudinal method is the lifeblood of developmental science. Unfortunately, the number of published longitudinal dMRI studies of children and adolescents is woefully small. More longitudinal studies with large samples and multiple time-points per participant, also incorporating methodological improvements discussed earlier in this review, are sorely needed. Greater sample sizes are especially important to give sufficient statistical power to further explore sex differences in WM development during puberty, as well as individual differences in WM development and how these relate to genetic or early environmental influences, or developmental changes in cognition or symptomatology. As previously pointed out by others (Simmonds et al., 2014), studies with more within-individual measurements are needed to achieve enough sampling through the age-range to properly model individual complex non-linear change trajectories. Datasets with multiple time-points per participant can additionally also allow for more sophisticated longitudinal statistical analyses, such as latent class growth analysis.

Second, inconsistencies regarding the precise developmental trajectories, especially for AD, and also in reported sex differences, call into question the comparability of samples and/or methods used and highlight the need for replication and method studies. Promising approaches for addressing these issues include multisample longitudinal studies from diverse populations and scanners/sequences, either studies pooling data and statistically testing for the effects of such variables or studies repeating the same processing and statistical analyses across independent datasets as recently done for morpohometric structural neuroimaging data (Mills et al., 2016), and studies which in parallel analyze the same datasets using different methodological approaches (Snook et al., 2007).

Third, multimodal studies exploring the relationships between different imaging metrics are called for as they promise to increase the specificity of our interpretation of the measurements and their changes over time and eventually give a more comprehensive understanding of brain development. For instance, regional WM volumes and DTI indices in WM have been shown to be only weakly to moderately related, and these measures are thus believed to be differentially sensitive to tissue characteristics and to provide complimentary information (Fjell et al., 2008; Tamnes et al., 2010). Critically, longitudinal data from children and adolescents also indicate that representations of tract volume increases are not clearly associated with tract FA increases or MD decreases, supporting the conclusion that observed DTI parameter changes reflect microstructural development rather than gross anatomy (Lebel & Beaulieu, 2011). Importantly (but not surprisingly), this means that caution should be taken when comparing results from DTI and volumetric studies, and furthermore that multimodal studies are warranted when investigating developmental changes in WM. DTI approaches should also be combined with other likely more specific diffusion models, such as NODDI, as well as with measurements of anatomical structural covariance and functional connectivity to increase our understanding of developmental changes in brain connections and networks.

Last, in our opinion, a diversity of types of studies, including both large “population neuroscience” studies, allowing investigations of for instance influencing genetic and environmental factors and rare disorders, and smaller method-focused or hypothesis-driven studies, is needed to progress the field. Examples of the latter can include studies with temporally dense longitudinal imaging across multiple time-points which allow for exploration of rapid changes or fluctuations in DTI indices (see e.g., (Elvsashagen et al., 2015; Hofstetter et al., 2013)), for instance across important transition periods in life or in combination with specific interventions, and more detailed longitudinal tracking of the associations between rapidly occurring and individually age-variable changes in hormone levels in puberty and WM microstructure.

In conclusion, in addition to providing novel insights into brain development, dMRI holds promise as a useful tool for elucidating sex differences in WM development during puberty, the influence of genetic and environmental factors on brain maturation, the role WM development plays in cognitive and behavioral changes during childhood and adolescence, and aberrant WM development in clinical or at-risk groups. Future longitudinal studies combined with methodological advances might provide further insights into brain development in childhood and adolescence. Importantly, dMRI measures and their longitudinal reliability are however influenced by hardware, scanner software, acquisition parameters, processing and statistical analysis methods, and researchers investigating brain development need to be mindful of this when designing and performing longitudinal studies and when analyzing and interpreting the results.

